# Regulation of oligodendrocyte lineage development by VAMP2/3 in the developing mouse spinal cord

**DOI:** 10.1101/2022.09.10.507429

**Authors:** Christopher D. Fekete, Robert Z. Horning, Matan S. Doron, Akiko Nishiyama

## Abstract

In the developing and adult CNS, new oligodendrocytes (OLs) are generated from a population of cells known as oligodendrocyte precursor cells (OPCs). As they begin to differentiate, OPCs undergo a series of highly regulated changes to morphology, gene expression, and membrane organization. This stage represents a critical bottleneck in oligodendrogenesis, and the regulatory program that guides it is still not fully understood. Here we show that *in vivo* toxin-mediated cleavage of the vesicle associated SNARE proteins VAMP2/3 in the OL lineage of both male and female mice impairs the ability of early OLs to mature into functional, myelinating OLs. In the developing mouse spinal cord, many VAMP2/3-deficient OLs appeared to stall in the premyelinating, early OL stage, resulting in an overall loss of both myelin density and OL number. The Src kinase Fyn, a key regulator of oligodendrogenesis and myelination, is highly expressed among premyelinating OLs, but its expression decreases as OLs mature. We found that OLs lacking VAMP2/3 in the spinal cord white matter showed significantly higher expression of Fyn compared to neighboring control cells, potentially due to an extended premyelinating stage. Overall, our results show that functional VAMP2/3 in OL lineage cells is essential for proper myelin formation and plays a major role in controlling the maturation and terminal differentiation of premyelinating OLs.

**Significance Statement:** The production of mature oligodendrocytes (OLs) is essential for CNS myelination during development, myelin remodeling in adulthood, and remyelination following injury or in demyelinating disease. Prior to myelin sheath formation, newly formed OLs undergo a series of highly regulated changes during a stage of their development known as the premyelinating, or early OL stage. This stage acts as a critical checkpoint in OL development, and much is still unknown about the dynamic regulatory processes involved. In this study, we show that VAMP2/3, SNARE proteins involved in vesicular trafficking and secretion, play an essential role in regulating premyelinating OL development and are required for healthy myelination in the developing mouse spinal cord.

## Introduction

During development, the generation of new oligodendrocytes (OLs) is essential for proper CNS myelination. Once established, mature OLs appear to be exceptionably stable and long lived, persisting well into adulthood (Tripathi et al., 2017) and for decades in the human brain (Yeung et al., 2014). Despite the apparent stability of established OLs, oligodendrogenesis continues throughout adulthood, contributing to experience-dependent myelination and remodeling (Young et al., 2013; Hughes et al., 2018).

Both during development and in adulthood, OLs are derived from oligodendrocyte precursor cells (OPCs), a population of proliferative cells that persists throughout adult life (Kessaris et al., 2006; Nishiyama et al., 2016). During differentiation, OPCs must undergo a series of progressive changes in morphology, cell to cell interactions, and gene expression to achieve the specialized structure and function of mature, myelinating OLs. These steps are highly regulated and a significant portion of newly formed OLs fail to integrate and are lost (Barres et al., 1992; Trapp et al., 1997; Hughes et al., 2018; Sun et al., 2018). This stage of OL development is often referred to as the early OL or premyelinating OL stage and typically lasts for 2-3 days in mice (Hill et al., 2014; Hughes et al., 2018). In zebrafish, the formation of new myelin sheaths has been shown to occur over just a 5 hour period, further highlighting the need for strict temporal control over this process (Czopka et al., 2013).

One of the signaling molecules known to play an important role in regulating this process is the Src kinase Fyn, which is thought to be an important component of the axon-OL signaling axis required to initiate myelination (Umemori et al., 1994; Laursen et al., 2009; White and Krämer-Albers, 2014). Fyn has been shown to regulate early OL survival, morphology, and the timing of major myelin-associated gene expression (Colognato et al., 2004; White et al., 2008; Mitew et al., 2014; Matrone et al., 2020). Recent work has even shown that modulating Fyn activity during a critical time window of new myelin sheath formation can alter the number of myelin sheaths a given cell will form (Czopka et al., 2013).

Like all eukaryotic cells, OL lineage cells employ SNARE-mediated membrane fusion to enable secretion, vesicle and membrane trafficking, as well as targeting and sorting of membrane proteins, crucial functions in OL development (Feldmann et al., 2009). In the OL lineage, inhibition of vesicle-associated SNARE proteins VAMP3 or VAMP7 was shown to impair the trafficking and surface presentation of myelin proteolipid protein (PLP) in the OPC cell line, Oli-neu, and primary oligodendrocyte cultures (Feldmann et al., 2011). The transcription of myelin basic protein (MBP) has also been shown to be regulated by syntaxin 4, a target-membrane associated SNARE protein (Bijlard et al., 2015).

In order to better understand the role that SNARE mediated processes play in early OL lineage development, we utilized the iBot mouse line which expresses the light chain of Botulinum toxin type B (BoNT/B-LC) in a cre-dependent manner (Slezak et al., 2012). BoNT/B is a potent neurotoxin which blocks SNARE-mediated processes by cleaving VAMP1-3 (Schiavo et al., 1992; Humeau et al., 2000; Chen et al., 2008; Yamamoto et al., 2012). The light chain of BoNT/B contains the catalytically active portion of the toxin but lacks the heavy chain required for the toxin to enter into cells, restricting the toxin’s access to a targeted population of cells.

Here we crossed iBot mice with NG2cre mice to impair VAMP1-3 mediated exocytosis in OPCs and their progeny. OL lineage cells primarily express VAMP2-4 and VAMP7 (Feldmann et al., 2009; Zhang et al., 2014); however, since VAMP4 and VAMP7 are not targeted by BoNT/B, we primarily inhibited VAMP2/3 dependent processes. We found that cleavage of VAMP2/3 in this population resulted in spinal cord hypomyelination and a reduction in the number of mature OLs formed. Interestingly, we did not detect major changes in OPC generation, proliferation, or survival. Based on our results, VAMP2/3 function becomes critical during the final development of early OLs into mature OLs as loss of VAMP2/3 function impairs their ability to fully mature into stably integrated, myelinating OLs.

## Materials and Methods

### Animals and transgenic mouse lines

All animal procedures received approval by the Institutional Animal Care and Use Committee at the University of Connecticut. To cleave VAMP1-3 in oligodendrocyte lineage we crossed heterozygous iBot mice (iBot^tg/+^) with homozygous NG2cre mice (NG2cre^tg/tg^). Here we will refer to iBot^tg/+^NG2cre^tg/+^ animals as iBot^tg/+^ mice and control iBot^+/+^NG2cre^tg/+^ animals as iBot^+/+^ mice. The iBot mice express cre-dependent BoNT/B-LC followed by an IRES and EGFP reporter (Jackson Laboratory Stock No: 018056; B6.FVB-Tg(CAG-boNT/B,-EGFP)U75-56Fwp/J) (Slezak et al., 2012). The NG2cre mice used were originally generated using a BAC construct containing NG2creER^TM^ but were found to express cre in NG2+ cells independent of tamoxifen induction (Zhu et al., 2011). Both male and female offspring P13-14 were used for all experiments. iBot^tg/+^ mice were identified either by genotyping PCR or on the basis of their motor function with subsequent confirmation of GFP expression. Primer sequences used for genotyping can be found in Table 1. For EdU pulse chase experiments, animals were injected with 50mg/kg EdU (Cayman, 20518) three times with two hours between injections at P10 before being sacrificed at P13.

### qRT-PCR

For relative RNA quantification, total RNA from the spinal cords of three P13 iBot^tg/+^ and three P13 iBot^+/+^ mice were isolated using PureLink RNA Mini kit (InVitrogen), and DNA was removed by DNase I digestion. After the integrity of the extracted RNA was confirmed with an Agilent 4200 High Sensitivity RNA Screen Tape Assay (Center for Genome Innovation, University of Connecticut), 2 μg of total RNA from each sample was reverse transcribed using Superscript IV (InVitrogen) and a mixture of 20 μM random hexamer and 20 μM oligo(dT)20 primers. Subsequent qPCR was carried out on a CFX06 Real-Time PCR Detection System (Bio-Rad) using the primers listed in Table 1 at 300 nM (Integrated DNA Technologies) and SYBR Green (InVitrogen).

### SDS-PAGE and Western blot

For immunoblotting, spinal cords from P14 iBot^tg/+^ and iBot^+/+^ mice were either processed immediately or snap-frozen in liquid nitrogen and stored at −70°C until further processing. Protein was extracted via homogenization with a Dounce or plastic pestle homogenizer in 150-300 μl RIPA buffer containing 1× Halt Protease inhibitor cocktail (ThermoFisher) and 1x PhosSTOP (Roche or Sigma; VAMP3 and FYN Western Blots only), after which the homogenates were incubated on ice for 10-30 minutes. The lysate was then cleared by centrifugation at 13,000*g* for 10-15 minutes at 4°C. Protein concentration was measured using the *DC* Protein Assay kit (Bio-Rad), a detergent compatible modification of the Lowry method. Cleared lysates were heated for 10 minutes at 70°C in reducing sample bufferFor each sample, 50-100 μg of protein in 10-40 μl volume was electrophoresed through 8% or 12% Bolt Bis-Tris Plus gels (InVitrogen) in Bolt MOPS or MES SDS running buffer (ThermoFisher) and transferred to Immobilon-FL membrane (Millipore) in a transfer buffer consisting of 1.25 mM Bicine, 1.25 mM Bis-Tris, and 1 mM EDTA (pH7.2) at 100V for 1-1.5 hours in a chilled blotting cell. The membranes were blocked for 1 hour at room temperature in Odyssey PBS or TBS Blocking Buffer (LI-COR) and then incubated in primary antibodies (Table 2) diluted in blocking buffer containing 0.1% Tween20 overnight at 4°C. Following 4 washes in PBS or TBS containing 0.1% Tween20, the membranes were incubated in secondary antibodies conjugated with IRDye680 or IRDye800 (LI-COR) diluted in blocking buffer containing 0.1% Tween20 and 0.01% SDS for 1 hour at room temperature. After 4 washes in PBS or TBS with 0.1% Tween20, the membranes were imaged on an Odyssey Infrared Imaging System (LI-COR) using Odyssey Imaging Software (LI-COR). The intensity of the bands was quantified using Image Studio Lite Quantification Software (LI-COR) or using the FIJI distribution of ImageJ (Schindelin et al., 2012).

### Tissue processing and immunofluorescent labeling

To prepare the spinal cords used in this study, mice were perfused first with a PBS rinse followed by 4% paraformaldehyde with 0.1M L-lysine and 0.01M sodium metaperiodate in 0.1M phosphate buffer (PB). The tissue was then harvested and postfixed for 1-3 hours in the same fixative at 4°C before being washed four times in 0.2M PB, leaving the tissue overnight in the fourth wash at 4°C. The tissue was then moved to 30% sucrose in 0.1M PB for cryoprotection. After cryoprotection, the tissue was embedded in frozen O.C.T. compound (Sakura Tissue Tek or Fisher brand O.C.T). 20μm sections were cut onto Superfrost Plus slides (Fisherbrand) using a Leica CM3050S cryostat. 40μm free floating sections were cut using a freezing microtome.

For immunofluorescent labeling, sections were washed three times in 10mM phosphate-buffered saline (PBS) and blocked with 5% normal serum (either donkey or goat) and 0.3% Triton X-100 in PBS. An additional 5% BSA was included when blocking prior to using bovine-raised secondary antibodies. After blocking, tissue sections were incubated with primary antibodies overnight at either 4°C or RT. Primary antibody concentrations can be found in Table 2. Following primary antibody incubation, sections were washed three to four times in PBS before incubation with fluorescently labeled secondary antibodies for 1 hour at RT. All antibody diluent consisted of 0-5% donkey or goat serum and 0.2-0.3% Triton X-100 in PBS. Species-specific secondary antibodies were acquired from either Jackson ImmunoResearch or Thermo Fisher and were raised in either donkey, goat, or bovine. Secondary antibodies were conjugated to either Alexa Fluor 488 (used at 1:1000), Alexa Fluor 546 (1:500), Cy3 (1:500), or Alexa Fluor 647 (1:200). After secondary antibody incubation, sections were washed three to four times in PBS before mounting in either Vectashield antifade mounting media with DAPI (Vector Laboratories) or Prolong Gold antifade mounting media with DAPI (Thermo Fisher).

For detection of EdU, sections were removed from the secondary antibody incubation, washed three times in PBS, and incubated for 30 minutes at RT in a homemade Click reaction mixture containing 4 mM CuSO_4_·5H_2_O (Sigma), 4 μM Alexa Fluor-647-conjugated azide (ThermoFisher), and 100 mM sodium ascorbate (Sigma) in Tris buffered saline pH 7.15. Following this, sections were washed in 3% BSA in PBS twice followed by one wash in PBS. Sections were then counterstained with Hoechst’s 33342 before being mounted using Vectashield without DAPI (Vector Laboratories).

For myelin staining, Fluoromyelin Red (Invitrogen) was used on cryostat sections from tissue perfused as described above that had been additionally subjected to an overnight post-fixation, prior to cryoprotection and embedding. Staining was performed following the manufacturer’s supplied protocol.

### Image acquisition and analysis

Images were primarily captured using a Nikon A1R Confocal with either a Plan Apo VC 60x/1.4 oil objective, a Plan Fluor 40x/1.3 oil objective, or a Plan Apo λ 20x/0.75 objective. Some images were collected using a Leica SP8 Confocal with either a Plan Apo HC 20x/0.75 or Plan Apo HC 40x/1.30 oil objective. Images for EdU pulse chase analysis and GSTπ labeling were captured on a Zeiss Axiovert 200M with attached Apotome using either a Plan Apochromat 20x/0.75 objective or Plan NeoFluar 40x/1.3 oil objective. Image analysis and manipulation was performed using the FIJI distribution of ImageJ (Schindelin et al., 2012).

For quantification of cell density and proportion, we used confocal or Apotome z-stacks collected from the dorsal column region. The boundaries of the dorsal column were outlined and used to calculate total volume of the counted region. The number of immunoreactive cells for each count was divided by the volume (in mm^3^) to calculate density. To calculate the percentage of PLP/DM20+ cells with premyelinating morphology, we collected confocal z-stacks from the dorsal horn spinal cord gray matter and scored cells as either myelinating or premyelinating based on the number and complexity of their processes as well as whether they appeared to contact and form sheaths around axons, using the CC1 antibody to Quaking 7 (QKI7) (Bin et al., 2016) to positively identify OLs. It is important to note that the different stages of OL development represent a continuous progression rather than clearly defined states; therefore, cells were scored based on their overall morphology, and some cells were excluded where it was not possible to definitively identify their developmental state.

To measure the fluorescence intensity of Fyn and p416 SFK in GFP+/QKI7+ and GFP-/QKI7+ OLs, we visually divided the dorsal column into three component white matter tracts: the corticospinal tract (cst), the cuneate fasciculus (cf), and the gracile fasciculus (gf). Using a Plan Fluor 40x/1.3 oil objective and a z-step size of 0.25μm, we collected one z-stack from each sub-region for each animal. We then selected 20 z-frames from each stack to create maximum projection images. Using the maximum projection images, we drew ROI’s around three GFP+/QKI7+ cells and three GFP-/QKI7+ cells in each image. To restrict analysis just to the outer border of each cell, a smaller ROI was created 10 pixels inside the first ROI. We then combined the two ROIs with the XOR operator to generate a donut-shaped final ROI. This final ROI was used to measure the fluorescence intensity of Fyn or p416 SFK on summed projection images constructed from the original 20 z-frames (each frame’s fluorescence intensity was summed into a single projection image). Additional details for specific images or analyses can be found in the figure legends.

### Experimental design and statistical analysis

In all cases, experiments were designed to compare either iBot^tg/+^ with iBot^+/+^ animals or GFP+ (BoNT/B-LC reporter) vs GFP-cells in iBot^tg/+^ animals. In one case, comparisons were made both among GFP+ and GFP-cells as well as among iBot-expressing and control animals (%ASPA+ OLs; Figure 4E). For statistical testing, all comparisons between two groups were conducted using Welch’s two sample t test with a significance cutoff of p < 0.05. For qPCR and Western Blot data, individual t tests were used to compare iBot-expressing and control animals for each gene/protein. For comparison of three groups (Figure 4E), a one-way ANOVA was performed followed by Tukey HSD post hoc testing. All statistical analysis and graphing were performed in R (R Core Team, 2021) using R studio (RStudio Team, 2021). The tidyverse package for R was used for data manipulation and graph generation (Wickham et al., 2019). Additional details for specific statistical tests are available in the figure legends.

## Results

### Expression of BoNT/B-LC in NG2+ cells and their progeny

To block the function of VAMP2/3 in the developing OL lineage, we crossed heterozygous iBot mice (iBot^tg/+^) expressing cre-dependent BoNT/B-LC and an IRES-linked EGFP reporter (Slezak et al., 2012), with homozygous NG2cre mice (NG2cre^tg/tg^; Figure 1A) (Zhu et al., 2011). In offspring heterozygous for the iBot transgene (referred to as iBot^tg/+^ mice), GFP expression was observed in OL lineage cells throughout the spinal cord in both gray and white matter. In the spinal cord dorsal column at P13, immunofluorescence staining for the OPC-specific marker PDGFRα (Figure 1B1) showed that 84.6 ± 3.7% of PDGFRα+ OPCs were GFP+, indicating that BoNT/B-LC was expressed in a majority of immature OPCs during development. To identify early and mature OLs, we labeled with the CC1 monoclonal antibody that detects Quaking 7 (QKI7; Figure 1B2) (Bin et al., 2016). Here we saw that only 46.4 ± 4.0% of QKI7+ OLs were GFP+, suggesting that a larger than expected fraction of OLs observed at this timepoint were derived from non-recombined OPCs. In order to assess the functional expression of BoNT/B in the iBot^tg/+^ mice we performed Western blotting for VAMP3 in spinal cord lysates from iBot^+/+^ and iBot^tg/+^ mice (Figure 1C). Normalizing the VAMP3 signal to GAPDH, we observed a significant reduction in the amount of intact VAMP3 in the iBot^tg/+^ samples (2.76×10^3^ ± 586.16 a.u.) compared to controls (8.04×10^3^ ± 498.40 a.u.; Figure 1D), verifying that BoNT/B-LC was both expressed and active in the iBot^tg/+^ mice.

**Figure 1:**
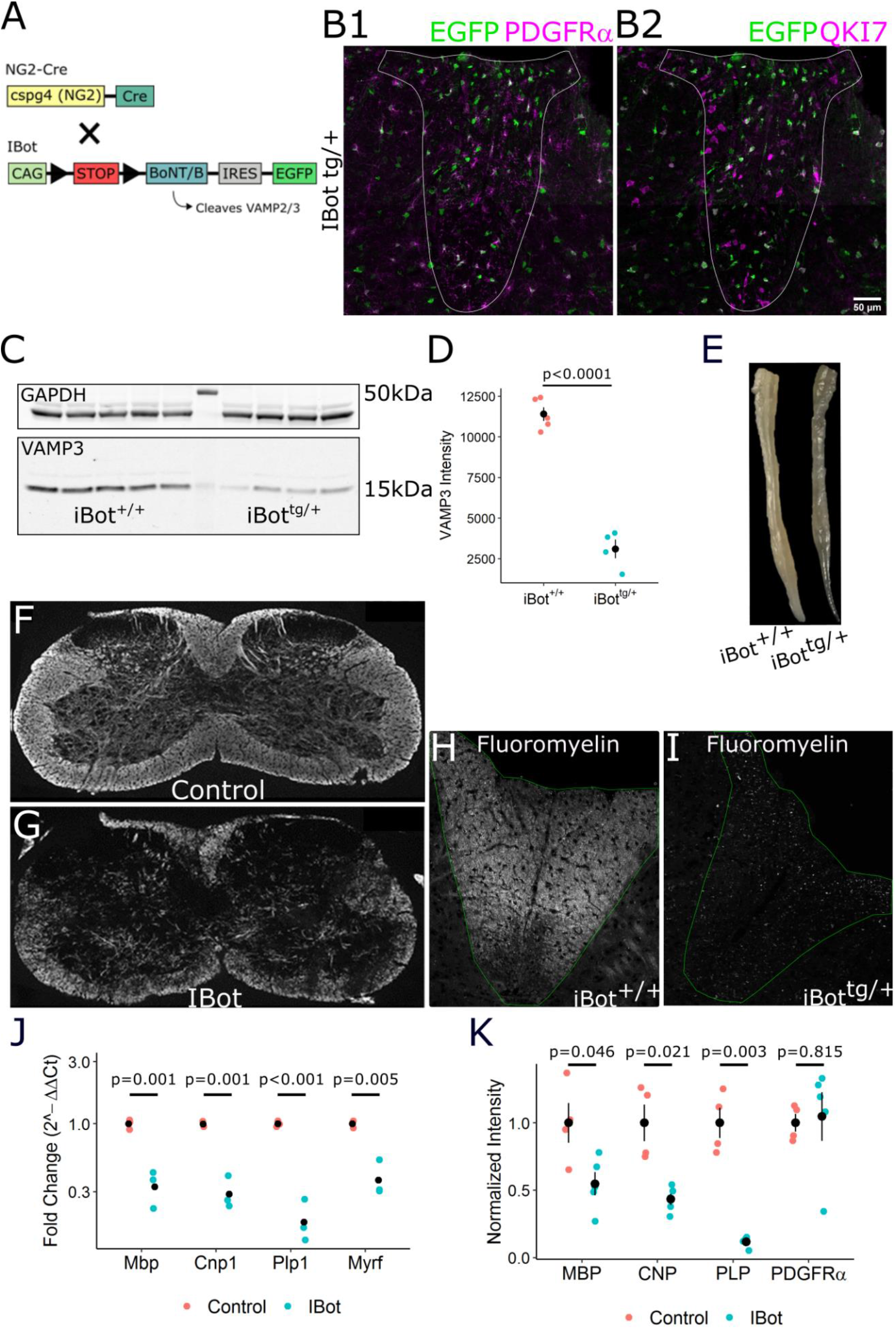
Developmental hypomyelination in iBot^tg/+^ spinal cords. **(A)** Schematic showing generation of iBot^tg/+^NG2cre^tg/+^ animals. **(B)** P13-14 iBot^tg/+^ dorsal column labeled for GFP and PDGFRα **(B1)** or QKI7 **(B2)** (scale bar = 50μm). **(C)** Western blot for VAMP3 (bottom) and GAPDH (top) from P14-15 iBot^+/+^ and iBot^tg/+^ spinal cord lysate **(D)** Quantification of VAMP3 levels normalized to GAPDH in iBot^+/+^ (8.04×10^3^ ± 498.40) and iBot^tg/+^ (2.76×10^3^ ± 586.16) spinal cords (Welch two sample t-test, t(5.83) = 11.7, p = 2.87×10^−5^, n = 5 iBot^+/+^ and 4 iBot^tg/+^). **(E)** Fixed spinal cords harvested from an iBot^+/+^ control animal (top) and an iBot^tg/+^ animal (bottom). **(F, G)** Cervical spinal cord sections labeled for MBP in iBot^+/+^ **(F)** and iBot^tg/+^ animals **(G). (H, I)** Fluoromyelin staining in the dorsal column of P13-14 iBot^+/+^ **(H)** and iBot^tg/+^ **(I)** mice. **(J)** qRT-PCR showing decreased expression of *Mbp* (3-fold decrease), *Cnp* (3.4-fold decrease), *Plp1* (5.4-fold decrease), and *Myrf* (2.6-fold decrease) in iBot^tg/+^ spinal cords (Welch two sample t test for each gene, n = 3 animals per genotype). Expression values are reported as fold change compared to controls plotted on a log10 scale **(K)** Western blot analysis showing decreased protein levels of MBP (1.8-fold decrease), CNP (2.3-fold decrease), and PLP (8.6-fold decrease) but not PDGFRα in iBot^tg/+^ animals compared to controls (Welch two sample t test for each protein, n = 4 animals per genotype). Band intensity shown is normalized to control values after normalization to the reference marker GAPDH. See extended data Figure 1-1 for video demonstrating motor impairment in an iBot^tg/+^ mouse.

### Blocking VAMP2/3-dependent exocytosis in the OL lineage caused severe hypomyelination in the developing mouse spinal cord

Between P10 and P13, iBot^tg/+^ mice began exhibiting motor defects which distinguished them from their control, iBot^+/+^ littermates. By ~P13, these behaviors included impaired limb function, movement difficulties, and difficulty righting themselves, and most of the iBot^tg/+^ mice ultimately died between P15-P17 (Video 1). Spinal cords harvested from iBot^tg/+^ mice frequently appeared more translucent than those from littermate controls, suggesting that the motor impairment may be due to a loss of myelin density (Figure 1E).

Immunofluorescent staining for MBP in spinal cord sections at P13 showed that iBot^tg/+^ animals had patchy, less dense myelin staining compared to controls (Figure 1F, 1G). Staining with Fluoromyelin in the dorsal column white matter further indicated a substantial reduction in myelin density in the iBot^tg/+^ (Figure 1I) spinal cord white matter compared to controls (Figure 1H). Consistent with this, qRT-PCR analysis from whole spinal cord isolate revealed decreased expression of major myelin genes *Mbp*, *Cnp1*, *Plp1*, and *Myrf* in iBot^tg/+^ animals relative to controls (Figure 1J). Western blot analysis of whole spinal cord homogenate similarly showed lower protein levels of MBP, CNP, and PLP but not PDGFRα (Figure 1K). Together, these results suggested a significant deficiency in either myelin generation or oligodendrocyte differentiation without a major impact on OPCs or their expression of PDGFRα, a key regulator of OPC proliferation and differentiation (Mitew et al., 2014).

### Loss of VAMP2/3 function in OPCs significantly reduced the density of QKI7+ OLs but not OPCs

In order to determine if the myelin deficiency seen in iBot^tg/+^ animals was due to a defect in oligodendrocyte differentiation, we measured the density of PDGFRα+ OPCs (Figure 2A, 2B) and QKI7+ OLs (Figure 2D, 2E) in the dorsal column white matter. At P13, we found no significant difference in the density of PDGFRα+ cells in iBot^tg/+^ animals (5.72×10^4^ ± 2.05×10^3^ cells/mm^3^) compared to controls (4.84×10^4^ ± 6.25×10^3^ cells/mm^3^; Figure 2C); however, the density of early and mature OLs labeled with QKI7 was significantly lower (1.37×10^5^ ± 1.47×10^4^ cells/mm^3^ in iBot^tg/+^ vs 2.34×10^5^ ± 1.89×10^4^ cells/mm^3^ in iBot^+/+^; Figure 2F). The relatively high percentage of PDGFRα+ cells expressing GFP (~85%, Figure 1C) and the similar density of PDGFRα+ cells between iBot^tg/+^ animals and controls (Figure 2C) indicated that this reduction in QKI7+ OLs was not secondary to a loss of OPCs. We therefore reasoned either that developing OLs were being lost after reaching the QKI7+ stage, or that differentiation of OPCs was blocked or impaired. As we did not observe an accumulation of OPCs in the iBot^tg/+^ animals, we hypothesized that OPCs were differentiating into QKI7+ OLs but being lost before reaching full maturity.

**Figure 2:**
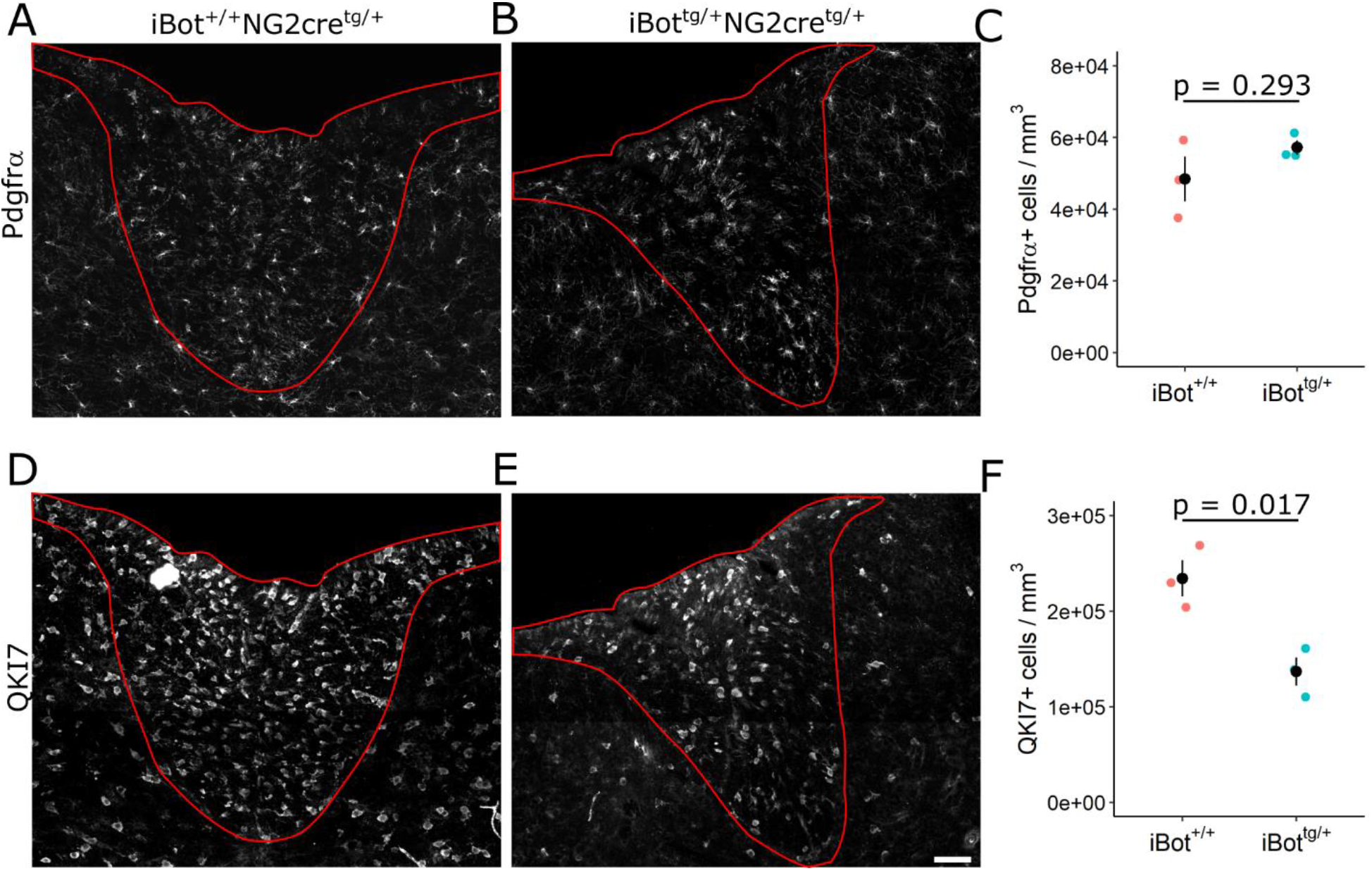
Density of PDGFRα+ OPCs and QKI7+ OLs in the dorsal column. **(A, B)** PDGFRα labeling in P13-14 iBot^+/+^ **(A)** and iBot^tg/+^ **(B)** dorsal column. **(C)** PDGFRα+ cell density in iBot^tg/+^ dorsal column (5.72 x 10^4^ ± 2.05×10^3^ cells/mm^3^) and control iBot^+/+^ dorsal column (4.84×10^4^ ± 6.25×10^3^ cells/mm^3^, Welch two sample t test, t(2.43) = −1.338, p =0.293, n = 3 animals for each genotype). **(D, E)** QKI7 labeling in P13-14 iBot^+/+^ **(D)** and iBot^tg/+^ **(E)** dorsal column. **(F)** QKI7+ cell density in iBot^tg/+^ dorsal column (1.37×10^5^± 1.45×10^4^ cells/mm^3^) and control iBot^+/+^ dorsal column (2.34×10^5^ ± 1.89×10^4^ cells/mm^3^, Welch two sample t test, t(3.77) = 4.074, p = 0.017, n = 3 animals for each genotype).

### OPCs lacking VAMP2/3 in iBot^tg/+^ mice initiated differentiation, but fewer mature OLs were formed

To assess whether the differentiation of OPCs into newly formed OLs was impaired in the iBot^tg/+^ animals, we performed EdU pulse chase labeling by injecting animals with EdU at P10 and sacrificing them P13 to label proliferating OPCs (Figure 3A). We then performed immunofluorescent labeling for QKI7 and EdU labeling in spinal cord sections of iBot^tg/+^ animals (Figure 3C) and controls (Figure 3B). The presence of EdU+/QKI7+/GFP+ cells indicated that iBot^tg/+^ OPCs did successfully differentiate to the QKI7+ early OL stage between P10 and P13 (Figure 3D, blue arrows). To determine if the overall number of new QKI7+ OLs produced during the three-day time course was affected, we measured the density of EdU+/QKI7+ cells in the dorsal column and found that there was not a significant difference between iBot^tg/+^ animals (1.67×10^4^ ± 4.97×10^3^ cells/mm^3^) and controls (1.82×10^4^ ± 6.16×10^3^ cells/mm^3^; Figure 3E) despite significantly fewer QKI7+ OLs overall (Figure 2F). This further indicated that the reduction in OL number seen in iBot^tg/+^ mice was not due to a failure of OPCs to generate new QKI7+ OLs, supporting the interpretation that VAMP2/3 deficiency primarily affects developing OLs after they have reached the QKI7+ early OL stage.

**Figure 3:**
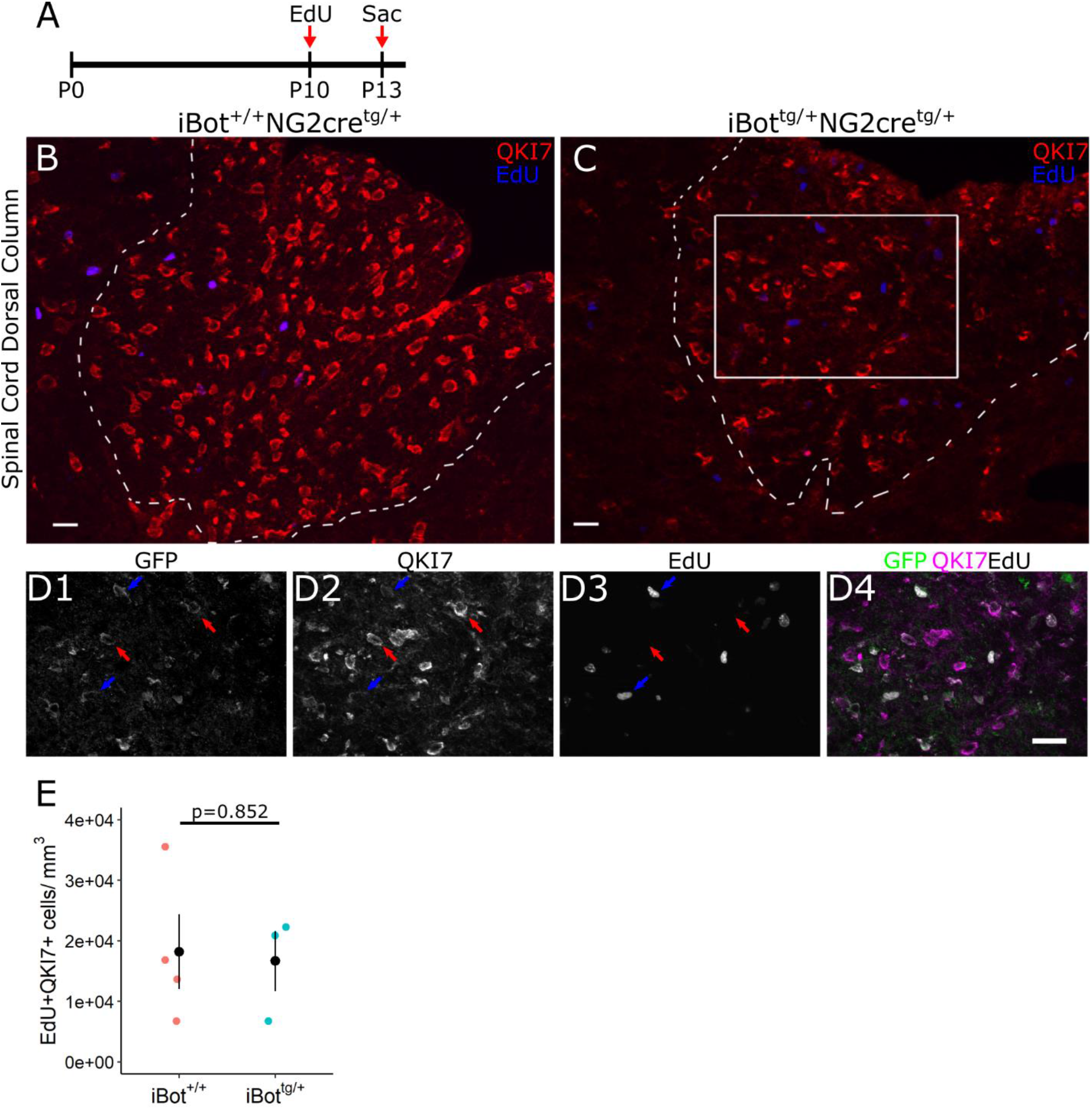
EdU pulse chase labeling of iBot^tg/+^ and control animals from P10-P13. **(A)** Schematic illustrating design of EdU pulse chase experiment. **(B, C)** Double label for QKI7 and EdU in P13 iBot^+/+^ **(B)** and iBot^tg/+^ **(C)** dorsal column. **(D)** Labeling for GFP, QKI7, and EdU from the area marked in **(C)** showing GFP+/QKI7+/EdU-OLs (red arrows) and GFP+/QKI7+/EdU+ OLs (blue arrows). **(E)** EdU+/QKI7+ cell density in iBot^+/+^ (1.82×10^4^ ± 6.16×10^3^ cells/mm^3^) and iBot^tg/+^ dorsal column (1.67×10^4^ ± 4.97×10^3^ cells/mm^3^, Welch two sample t test, t(5) = 0.196, p = 0.852, n = 4 iBot^+/+^ and 3 iBot^tg/+^ animals). Scale bars = 20 μm **(B, C)**, 25 μm (**D**).

We tested this possibility further using immunofluorescent labeling for QKI7 in combination with the mature OL marker glutathione S-transferase π (GSTπ) (Tansey and Cammer, 1991). QKI7+/GSTπ+ cells were considered to be mature OLs, while QKI7+/GSTπ-cells were considered newly formed, early OLs (Figure 4C). Consistent with the overall reduction in QKI7+ cell density (Figure 2F), iBot^tg/+^ animals showed fewer GSTπ+ OLs (Figure 4B) than controls (Figure 4A). Additionally, the proportion of QKI7+ cells that were GSTπ+ was significantly lower in iBot^tg/+^ animals (31.3 ± 3.6%) compared to controls (56.6 ± 1.0%; Figure 4D) suggesting that, despite reaching the QKI7+ stage, fewer early OLs were successfully reaching full maturity in the iBot^tg/+^ animals.

**Figure 4:**
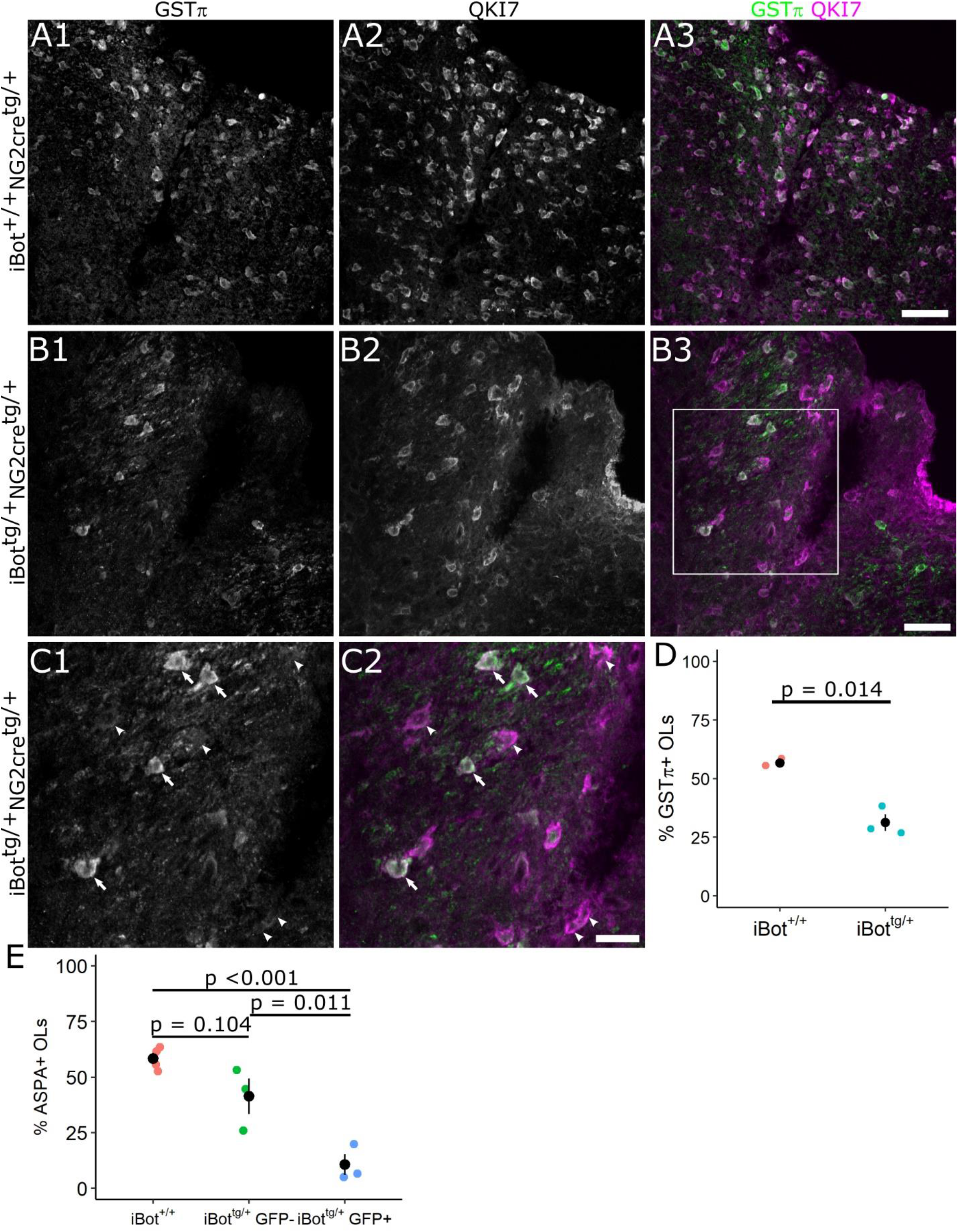
Percentage of OLs labeled by mature OL markers in P13-14 dorsal column. **(A, B)** Labeling for GSTπ and QKI7 at P13 in iBot^+/+^ **(A)** and iBot^tg/+^ dorsal column **(B)**. **(C)** Higher magnification view of the area marked in **(B)** showing QKI7+/GSTπ-early OLs (arrowheads) and QKI7+/GSTπ+ mature OLs (arrows). **(D)** Percentage of QKI7+ OLs also labeled by GSTπ in iBot^tg/+^ (31.3 ± 3.6%) and iBot^+/+^ dorsal column (56.6 ± 1.0%, Welch two sample t test, t(2.33) = 6.856, p = 0.014, n = 3 animals for each genotype). **(E)** Percentage of QKI7+ OLs that were ASPA+ at P13-14 in iBot^+/+^ dorsal column (58.4 ± 2.5%), among GFP-cells in iBot^tg/+^ dorsal column (41.3 ± 8.0%), and among GFP+ cells in iBot^tg/+^ dorsal column (10.6 ± 4.7%, ANOVA, F(2, 7) = 23.07, p = 8.30×10^−4^, Tukey HSD post hoc tests, p = 6.55×10^−4^ for iBot^+/+^:iBot^tg/+^GFP+, p = 0.011 for iBot^tg/+^GFP-:iBot^tg/+^GFP+, p = 0.104 for iBot^+/+^ to iBot^tg/+^GFP-). Scale bars = 50 μm **(A, B)**, 25 μm **(C)**.

Using another marker for mature OLs, aspartoacylase (ASPA) (Pan et al., 2020), we next tested whether the failure to reach a fully mature stage was restricted to those cells expressing BoNT/B-LC, identified as GFP+ cells. In control animals, we found that the proportion of QKI7+ cells that were also ASPA+ (58.4 ± 2.5%; Figure 4E) was similar to that seen with GSTπ (56.6 ± 1.0%; Figure 4D), confirming that the timing of expression for these two markers is similar in OL lineage development. In iBot^tg/+^ animals, the proportion of GFP+/QKI7+ cells labeled by ASPA was significantly lower (10.6 ± 4.7%) compared to the proportion of QKI7+ cells that were ASPA+ in control animals (58.4 ± 2.5%) and to the proportion of GFP-/QKI7+ cells that were ASPA+ sampled from iBot^tg/+^ animals (41.3 ± 8.0%; Figure 4E). This confirmed that fewer early OLs were becoming fully mature in the iBot^tg/+^ animals. In addition, this result suggested that VAMP2/3 deficiency impairs OL lineage development cell autonomously, as the proportion of GFP-OLs expressing ASPA in iBot^tg/+^ mice was not significantly reduced compared to controls.

In order to better determine the stage of differentiation at which iBot^tg/+^ OLs are affected, we labeled for NKX2.2, a transcription factor involved in the regulation of OL differentiation and maturation that is upregulated in OPCs just prior to the onset of differentiation and then rapidly downregulated as they approach the final stages of differentiation (Qi et al., 2001; Watanabe et al., 2004). Following a period of downregulation in newly-mature OLs, however, it has been shown that NKX2.2 is eventually re-expressed at lower levels (Cai et al., 2010; see Figure 10 for a diagram summarizing gene expression timing). Consistent with this, NKX2.2 labeling in the dorsal column of iBot^+/+^ spinal cords showed a mixture of both highly-expressing NKX2.2+ cells, corresponding to actively differentiating OLs (referred to as NKX2.2^high^ OLs), and weakly-expressing NKX2.2+ cells, corresponding to mature OLs that have reacquired NKX2.2 expression (referred to as NKX2.2^low^ OLs; Figure 5A). In comparison, in the dorsal column of iBot^tg/+^ mice we observed an apparent reduction in the overall density of NKX2.2+ cells as well as a relative lack of weakly-expressing NKX2.2^low^ cells (Figure 5B). When we quantified the overall density of NKX2.2+/QKI7+ OLs, using QKI7 to exclude NKX2.2+ OPCs, we found that the density was significantly reduced in the iBot^tg/+^ mice (2.08×10^4^ ± 2.30×10^3^ cells/mm^3^) relative to controls (3.93×10^4^ ± 1.80×10^3^ cells/mm^3^; Figure 5C). However, when we similarly quantified only the NKX2.2^high^/QKI7+ OLs we found that the density was not significantly affected in iBot^tg/+^ mice (1.33×10^4^ ± 2.42×10^3^ cells/mm^3^) relative to controls (9.29×10^3^ ± 263.90 cells/mm^3^; Figure 5D). This suggested that the number of actively differentiating OLs was unchanged but that the number of NKX2.2^low^ mature OLs was reduced in the iBot^tg/+^ mice, further supporting the interpretation that iBot^tg/+^ OLs proceed successfully through the initial stages of differentiation but either stall or undergo cell death prior to completing the maturation processes.

**Figure 5:**
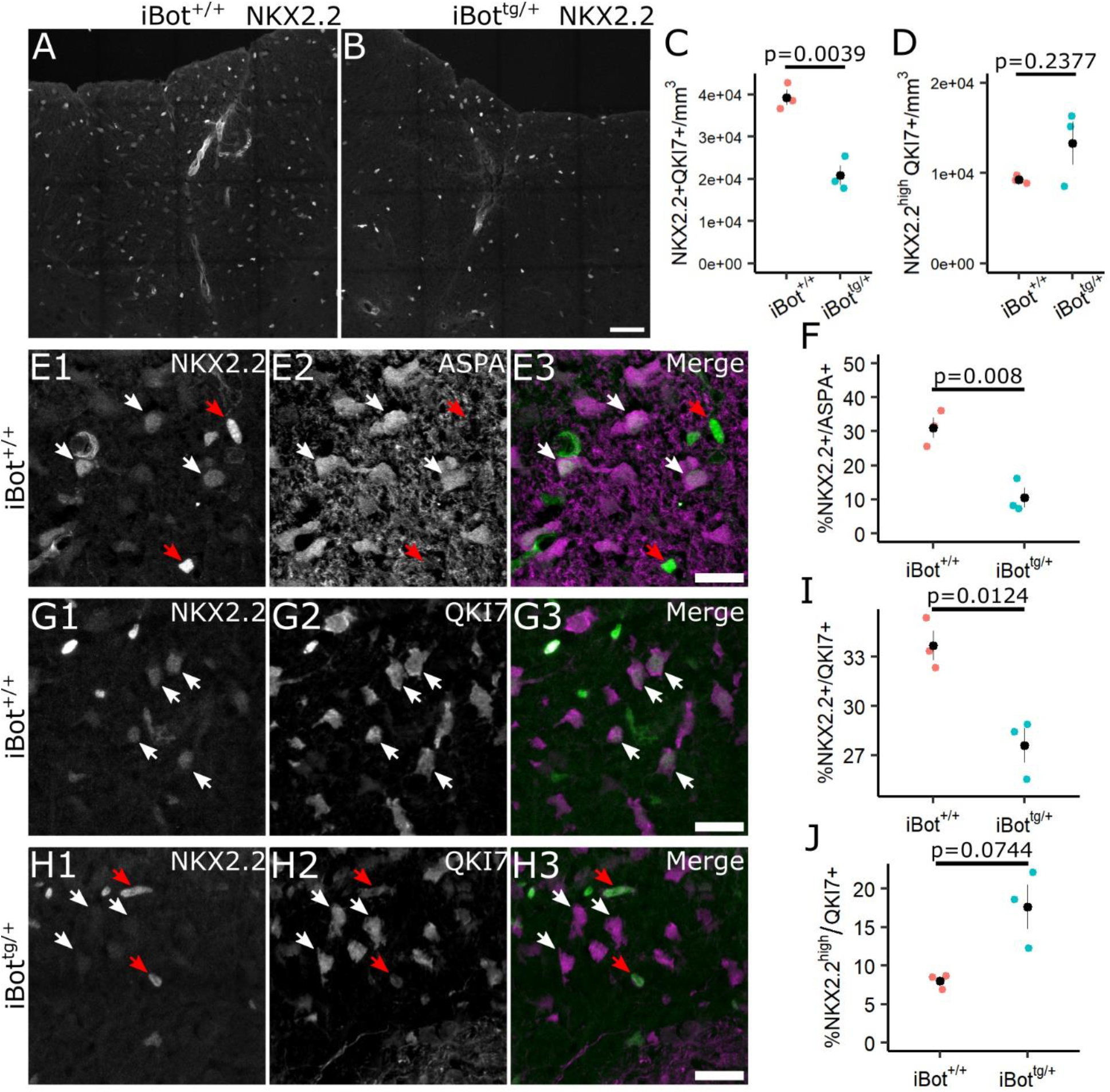
Expression of NKX2.2 among QKI7+ and ASPA+ OLs in iBot^+/+^ and iBot^tg/+^ dorsal column. **(A, B)** NKX2.2 labeling in spinal cord sections from P13-14 iBot^+/+^ **(A)** and iBot^tg/+^ **(B)** mice. **(C)** Density of NKX2.2+/QKI7+ cells in dorsal column of iBot^+/+^ (3.93×10^4^ ± 1.80×10^3^ cells/mm^3^) and iBot^tg/+^ mice (2.08×10^4^ ± 2.30×10^3^ cells/mm^3^; t(3.79) = 6.31, p = 0.0039). **(D)** Density of NKX2.2^high^/QKI7+ in dorsal column of iBot^+/+^ (9.29×10^3^ ± 263.90 cells/mm^3^) and iBot^tg/+^ mice (1.33×10^4^ ± 2.42×10^3^ cells/mm^3^; t(2.05) = −1.65, p = 0.2377). **(E)** Labeling for NKX2.2 and ASPA in the dorsal column of iBot^+/+^ control mice showing NKX2.2^high^/ASPA-cells (red arrows) and NKX2.2^low^/ASPA+ cells (white arrows). **(F)** Quantification of the percentage of ASPA+ OLs with detectable nuclear NKX2.2 expression in iBot^+/+^ (31.06 ± 3.05%) and iBot^tg/+^ (10.53 ± 2.83%) dorsal column (t(3.98) = 4.93, p = 0.008). **(G)** Immunofluorescent labeling for NKX2.2 and QKI7 in iBot^+/+^ dorsal column showing NKX2.2^low^ OLs (white arrows). **(H)** Labeling for NKX2.2 and QKI7 in iBot^tg/+^ dorsal column showing NKX2.2^high^ (red arrows) and NKX2.2-OLs (white arrows). **(I)** Quantification of the percentage of NKX2.2+ QKI7+ OLs in iBot^+/+^ (33.68 ± 0.90%) and iBot^tg/+^ dorsal column (27.60 ± 1.05%; t(3.90) = 4.40, p = 0.0124). **(J)** Percentage of NKX2.2^high^/QKI7+ OLs in iBot^+/+^ (8.01 ± 0.55%) and iBot^tg/+^ dorsal column (17.64 ± 2.88%; t(2.15) = −3.28, p = 0.0744). All statistical tests done using Welch two sample t-test with n = 3 animals for each genotype. Scale bars = 50μm **(A, B)**, 20μm **(E, G, *H*)**.

To confirm that the NKX2.2^low^ cells observed were fully differentiated OLs, we double labeled for NKX2.2 and ASPA as a marker for mature OLs. In iBot^+/+^ control mice, we found that many of the NKX2.2^low^ cells were also ASPA+, confirming they were fully mature OLs, while the NKX2.2^high^ cells were almost entirely ASPA-, indicating they were either immature, actively differentiating OLs or committed OPCs (Figure 5E). When we quantified the percentage of ASPA+ OLs that co-expressed NKX2.2 we found that it was significantly lower in the iBot^tg/+^ mice (10.53 ±2.83%) compared to controls (31.06 ± 3.05%), suggesting a reduced accumulation of mature OLs in the developing iBot^tg/+^ spinal cord (Figure 5F). Similarly, using double labeling for QKI7 and NKX2.2 in iBot^+/+^ and iBot^tg/+^ dorsal column (Figure 5G, 5H), we found that the percentage of QKI7+ OLs co-expressing NKX2.2 was also significantly reduced in iBot^tg/+^ mice (27.60 ± 1.05% vs 33.68 ± 0.90% in control mice; Figure 5I). As ASPA+ OL are also QKI7+, the milder reduction in the proportion of NKX2.2+ OLs among total QKI7+ OLs likely also reflects a reduced or slowed developmental accumulation of mature OLs reacquiring NKX2.2 expression. Notably, however, the percentage of QKI7+ OLs that were NKX2.2^high^ was not reduced, instead showing a trend towards an increased percentage of NKX2.2^high^/QKI7+ OLs in the iBot^tg/+^ mice (Figure 5J), suggesting that the iBot^tg/+^ OLs may undergo cell death, removing them from the total QKI7+ population, following the initial downregulation of NKX2.2 in the final stages of differentiation but prior to its re-expression in fully mature OLs.

### Survival of OLs appeared reduced in the dorsal column of iBot^tg/+^ animals

To further investigate the fate of differentiating iBot^tg/+^ OLs, we performed immunofluorescent staining for cleaved caspase 3 (cCasp3), a marker for cells undergoing apoptosis, to determine if their survival was impaired by loss of VAMP2/3 function. At P13, we found that the density of cCasp3+ cells in the dorsal column was significantly higher in iBot^tg/+^ mice (4.45×10^3^ ± 992.8 cells/mm^3^) compared to iBot^+/+^ control mice (1.64×10^3^ ± 429.9 cells/mm^3^; Figure 6C). Though it is worth nothing that few cCasp3+ cells were detected overall in either iBot^tg/+^ or control animals (range of 0-9 cCasp3+ cells per dorsal column sampled; Figure 6A,6B). To assess whether the cCasp3+ cells in iBot^tg/+^ animals belonged to the OL lineage, we then labeled for cCasp3 and for Olig2, a marker expressed throughout the OL lineage (Figure 6A,6B). We found that there was a trend toward increased cCasp3+/Olig2+ cell density in iBot^tg/+^ mice (3.89×10^3^ ± 1.59×10^3^ cells/mm^3^) compared to controls (868.8 ± 369.6 cells/mm^3^; Figure 6D); however, this difference was not statistically significant. Preliminary TUNEL staining performed at P8 similarly suggested enhanced cell death in the iBot^tg/+^ mice (data not shown).

**Figure 6:**
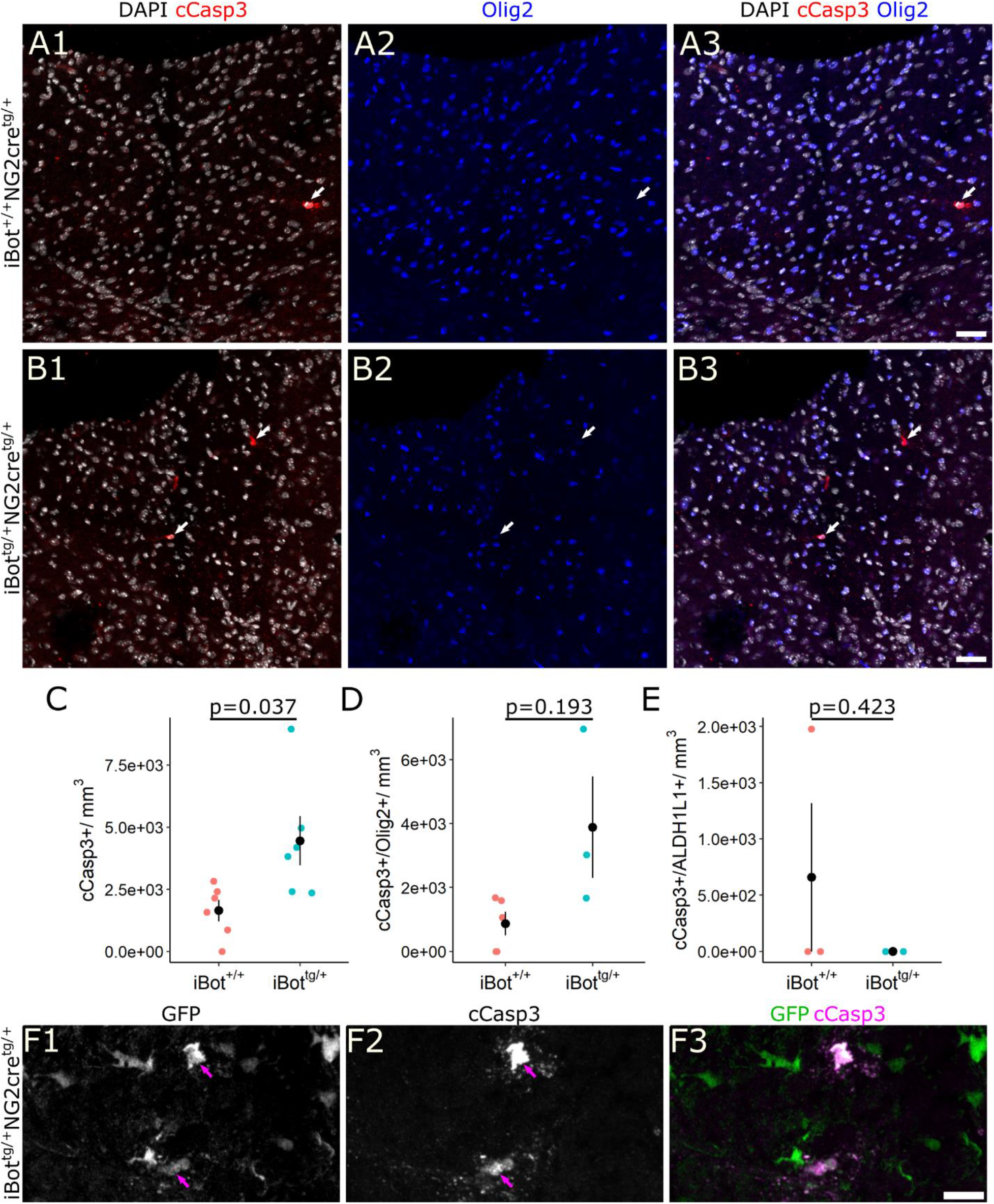
Cleaved Caspase 3 labeling for apoptotic cells in P13 dorsal column. **(A, B)** Double label for cCasp3 and Olig2 with DAPI in dorsal column of P13 iBot^+/+^ **(A)** and iBot^tg/+^ mice **(B)** showing cCasp3+ apoptotic cells (arrows). **(C)** cCasp3+ cell density in the dorsal column of P13 iBot^+/+^ mice (1.64×10^3^ ± 429.9 cells/mm^3^) and iBot^tg/+^ mice (4.45×10^3^ ± 992.8 cells/mm^3^, Welch two sample t test, t(6.8) = −2.60, p = 0.036, n = 6 iBot^+/+^ and 6 iBot^tg/+^ animals). **(D)** Density of double labeled cCasp3+/Olig2+ cells in the dorsal column of P13 iBot^+/+^ mice (868.8 ± 369.6 cells/mm^3^) and iBot^tg/+^ mice (3.89×10^3^ ± 1.59×10^3^ cells/mm^3^, Welch two sample t test, t(2.22) = -1.852, p = 0.193, n = 5 iBot^+/+^ and 3 iBot ^tg/+^ animals). **(E)** Density of double labeled cCasp3+/ALDH1L1+ cells in the dorsal column of P13 iBot^+/+^ mice (659.4 ± 659.4 cells/mm^3^) and iBot^tg/+^ mice (0 cells/mm^3^, Welch two sample t test, t(2) = 1, p = 0.4226, n = 3 animals for each genotype). **(F)** cCasp3+/GFP+ double labeled cells (arrows) in P13 iBot^tg/+^ dorsal column. Scale bars = 50 μm **(A, B)**, 10 μm **(F)**.

In order to assess the contribution of astrocytes undergoing apoptosis to the overall cCasp3+ cell density, we also performed immunofluorescent labeling with cCasp3 and the astrocyte marker ALDH1L1. The lack of cCasp3+/ALDH1L1+ astrocytes in iBot^tg/+^ animals indicated that apoptotic astrocytes did not contribute significantly to the overall cCasp3+ cell density (Figure 6E). Similarly, when we labeled with cCasp3 and the microglial marker Iba1, we detected no cCasp3+/Iba1+ microglia in the dorsal columns of iBot^tg/+^ animals (not shown). Comparison of cCasp3 staining with GFP expression in iBot^tg/+^ animals showed that 75.7% of cCasp3+ cells were also GFP+, suggesting that the majority of cCasp3+ cells were derived from NG2+ progenitors (Figure 6F). Taken together, these results suggest that the iBot^tg/+^ OLs that fail to reach maturity eventually undergo cell death resulting in the reduced density of QKI7+ OLs seen in the iBot^tg/+^ mice.

### At P13, fewer iBot^tg/+^ OLs display morphological features of mature, myelinating OLs

In the early OL phase, developing OLs begin expressing *Qki* and initially adopt a morphology that is distinct from more mature, myelinating OLs. OLs with a typical premyelinating morphology are characterized by numerous, highly-branched processes that extend radially from the soma. In contrast, myelinating OLs have fewer, less complex processes which extend toward and contact axons to form myelin sheaths (Butt et al., 1997).

To assess whether iBot^tg/+^ OLs underwent the stereotypical transition from premyelinating to myelinating morphology despite lacking the expression profile of fully mature OLs, we used an anti-PLP antibody, referred to by its clone name AA3 (Yamamura et al., 1991), that reacts with both PLP and the DM20 splice variant to compare the morphological characteristics of GFP+ and GFP-OLs in iBot^tg/+^ spinal cords. Here we examined OLs in the dorsal horn gray matter region where myelinated fibers are sparser, allowing us to visualize the morphology of individual cells more easily. Using QKI7 to identify both early and mature OLs in iBot^tg/+^ animals, we observed that GFP+/QKI7+ cells (OLs expressing BoNT/B-LC) could be found with both typical premyelinating morphology (Figure 7A) as well as more mature morphology indicative of myelination (Figure 7B) and comparable to GFP-/QKI7+ control cells (Figure 7C). This suggested that functional loss of VAMP2/3 did not uniformly block morphological maturation of iBot^tg/+^ OLs; however, we also found that a significantly higher percentage of GFP+/QKI7+ OLs had a premyelinating morphology (46.4 ± 6.9%) when compared to GFP-/QKI7+ OLs from the same animals (0%; Figure 6D). Consistent with this, iBot^tg/+^ animals appeared to have an expanded population of premyelinating OLs in the dorsal horn compared to controls (Figure 6E, 6F). Given that the number of newly formed QKI7+ OLs was unaffected in iBot^tg/+^ animals (Figure 3E, 5D), the increased prevalence of premyelinating cells among GFP+ OLs suggested that these cells may undergo an extended premyelinating stage, which could contribute to their ultimate failure to reach full maturity.

**Figure 7:**
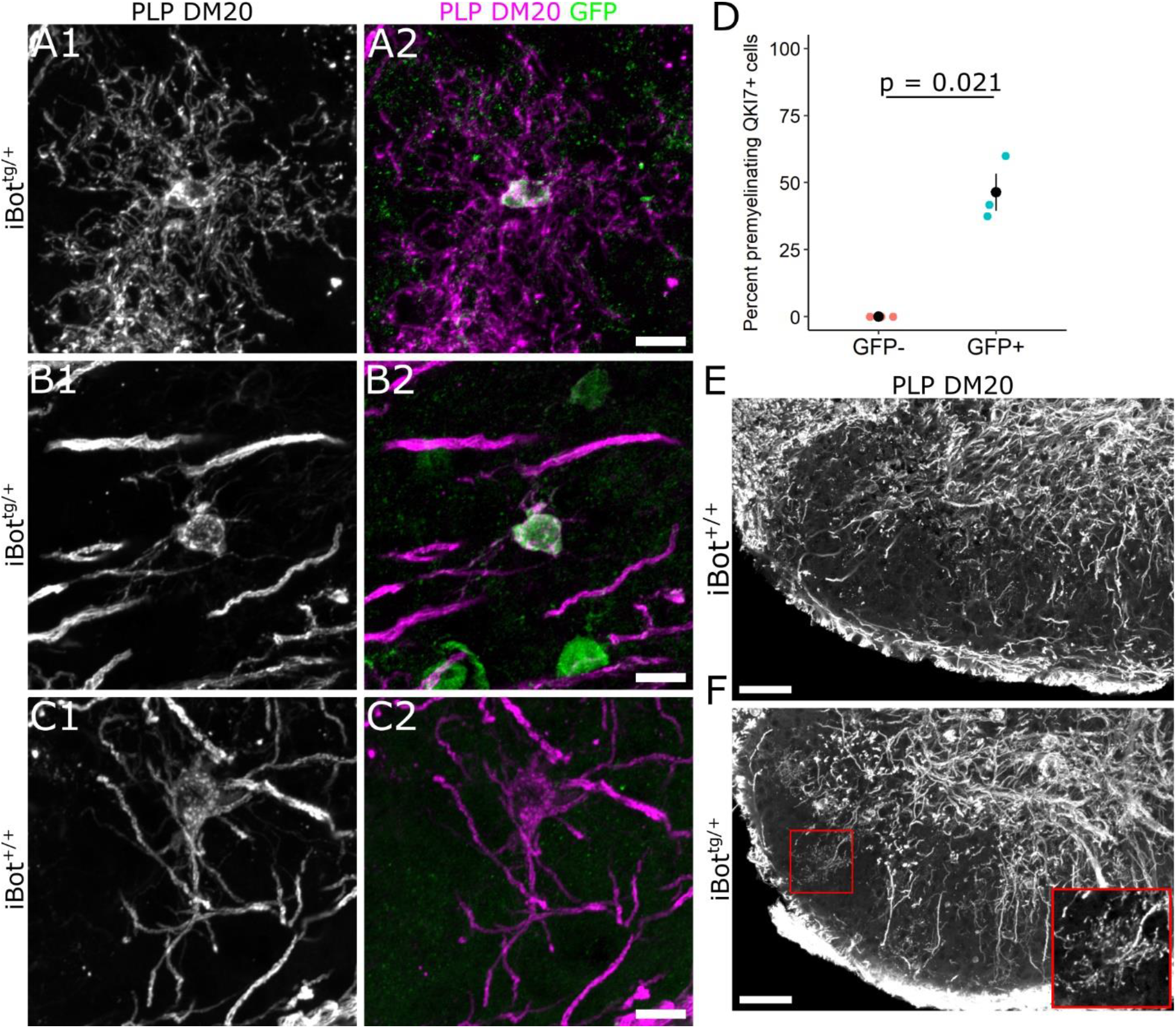
Morphological features of iBot^tg/+^ and iBot^+/+^ OLs in the dorsal horn at P13. **(A-C)** Double label for PLP/DM20 and GFP in iBot^tg/+^ **(A, B)** and iBot^+/+^ mice **(C)** showing a premyelinating OL **(A)** or myelinating OLs **(B, C)** in the dorsal horn at P13. **(D)** Percentage of QKI7+ OLs scored as having a premyelinating morphology among GFP-OLs (0%) and GFP+ OLs (46.4 ± 6.9%, Welch two sample t test, t(2) = −6.712, p = 0.021, n = 3 iBot^tg/+^ animals, cells were scored from two different images taken in the dorsal horn of each animal). **(E, F)** Low magnification images of PLP/DM20 labeling in the dorsal horn of iBot^+/+^ **(E)** and iBot^tg/+^ **(F)** mice at P13. Inset shows a magnified view of the premyelinating OL from the region outlined in red **(F)**. Gamma was adjusted to 0.5 **(E, F)** to better visualize premyelinating OLs. Scale bars = 10 μm **(A-C)**, 50 μm **(E, F)**.

### Early OLs lacking functional VAMP2/3 exhibited significantly higher levels of Src Kinase Fyn

We next wanted to investigate key signaling pathways known to regulate OL maturation that could be affected in the iBot^tg/+^ mice. As the differentiating iBot^tg/+^ OLs appeared to stall in the premyelinating stage around the onset of myelination, we looked for signaling factors that would be actively contributing to the regulation of this process. The non-receptor tyrosine kinase Fyn, a member of the Src family of kinases, has been shown to regulate OL lineage cell survival and maturation and is thought to be particularly important for mediating axon-OL signaling (Laursen et al., 2009; White and Krämer-Albers, 2014). In addition, recent scRNA-Seq data from OL lineage cells has shown that *Fyn* expression in the OL lineage is highly enriched in the late OPC/early OL stage and begins to decline as expression of major myelin associated genes like *Mag* and *Mog* increases (Marques et al., 2016; Floriddia et al., 2020). Immunofluorescent labeling for Fyn in QKI7+ OLs in the dorsal column of iBot^tg/+^ mice indicated significantly higher levels of Fyn associated with somas of GFP+/QKI7+cells (1.41×10^4^ ± 819.3 a.u.) compared to GFP-/QKI7+ controls (8.61×10^3^ ± 495.1 a.u.; Figure 8A, 8B). This suggested that the iBot^tg/+^ OLs may exhibit abnormally high levels of Fyn, possibly due to an accumulation of Fyn protein because of an extended premyelinating stage.

**Figure 8:**
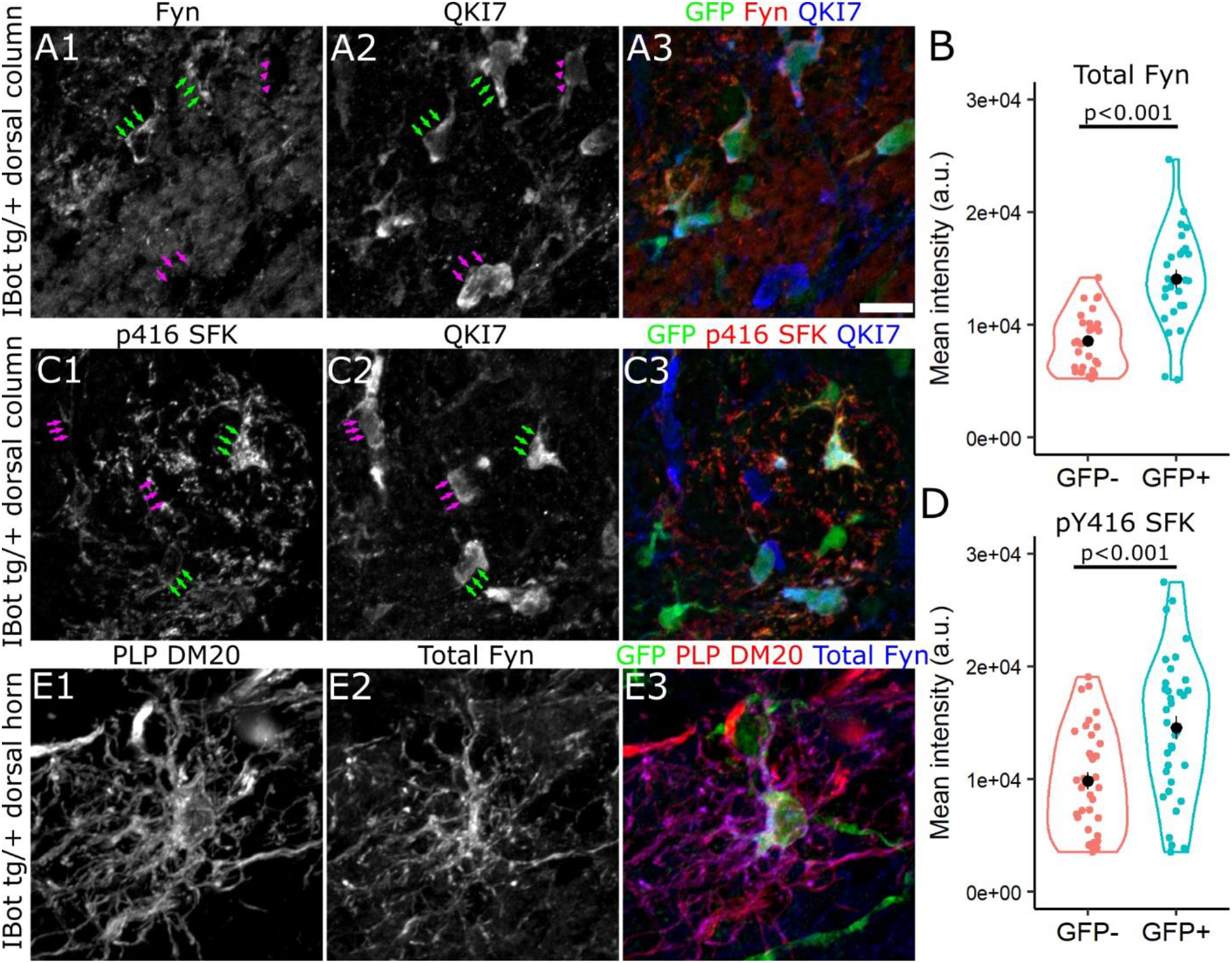
Fyn is highly expressed throughout the soma and processes of iBot^tg/+^ OLs. **(A, C)** Labeling for QKI7, GFP and either Fyn **(A)** or p416 SFK **(C)** around the border of iBot^tg/+^ (GFP+, green arrows) and iBot^+/+^ OLs (GFP-, magenta arrows) in iBot^tg/+^ corticospinal tract. **(B)** Average fluorescence intensity of Fyn staining along the edge of GFP-(8.61×10^3^ ± 495.1 a.u.) and GFP+ (1.41×10^4^ ± 819.3 a.u.) OLs in the dorsal column of iBot^tg/+^ animals (Welch two sample t test, t(42.76) = −5.70, p = 1.03×10^−6^, n = 27 cells per condition from 3 iBot^tg/+^ animals). **(D)** Average fluorescence intensity of p416 SFK staining along the edge of GFP-(9.83×10^3^ ± 757 a.u.) and GFP+ OLs (1.45×10^4^ ± 1.03×10^3^ a.u., Welch two sample t test, t(64.13) = −3.679, p = 4.81×10^−4^, n = 36 cells per condition from 4 iBot^tg/+^ animals). **(E)** GFP+ premyelinating OL labeled for PLP/DM20, Fyn, and GFP in P13 iBot^tg/+^ dorsal horn. Scale bar = 10 μm **(A, C)**.

Given that Fyn activity generally promotes maturation and myelination in early OLs, the possibility that iBot^tg/+^ OLs, which fail to reach maturity, showed enhanced levels of Fyn expression was surprising. Therefore, we also wanted to determine if the elevated levels of Fyn corresponded to functionally active Fyn as well. To test this, we performed similar immunofluorescent labeling with a phospho-specific antibody directed against Src family kinases phosphorylated at Y416 (pY416 SFK; Figure 8C). Fyn is the predominant Src family kinase expressed in the OL lineage and is activated by phosphorylation at pY416 (Matrone et al., 2020). Using QKI7 as a marker for developing and mature OLs, we compared the intensity of pY416 SFK staining around the somas of GFP- and GFP+ OLs in the dorsal column of iBot^tg/+^ animals and found that, similar to total Fyn, GFP+ OLs had significantly stronger pY416 SFK staining (1.45×10^4^ ± 1.03×10^3^ a.u.) compared to control cells (9.83×10^3^ ± 757 a.u.; Figure 8D).

As it was difficult to assess the distribution of Fyn within the processes of individual QKI7 labeled OLs in the dorsal column, we also wanted to address the possibility that Fyn may accumulate in the somas of iBot^tg/+^ OLs due to a VAMP2/3 dependent trafficking defect. To do this, we examined iBot^tg/+^ OLs in the dorsal gray matter using PLP-DM20 labeling to better visualize the morphology of individual cells. Here we saw strong Fyn staining not just in the soma and proximal processes, but throughout the more distal processes as well (Figure 8E), indicating that its transport into the processes was not defective in the iBot^tg/+^ OLs.

### Fyn expression is concentrated in NKX2.2-/GFP+ OLs in iBot^tg/+^ mice

To further investigate the differentiation status of the stalled iBot^tg/+^ OLs and clarify the relationship between Fyn kinase expression and the maturation defect observed, we performed double labeling for total Fyn and NKX2.2 in the dorsal gray matter of iBot^+/+^ and iBot^tg/+^ spinal cords. Compared to the dorsal gray matter of control mice, Fyn staining in iBot^tg/+^ mice appeared concentrated within relatively large, GFP+/NKX2.2-cells (Figure 9A, 9B). Their lack of NKX2.2 expression indicated that they were not actively differentiating OLs, but that they had also not reached the fully mature NKX2.2 re-expressing stage. A similar distribution of Fyn could also be observed in the dorsal cst white matter tract of iBot^tg/+^ spinal cords (Figure 9C).

**Figure 9:**
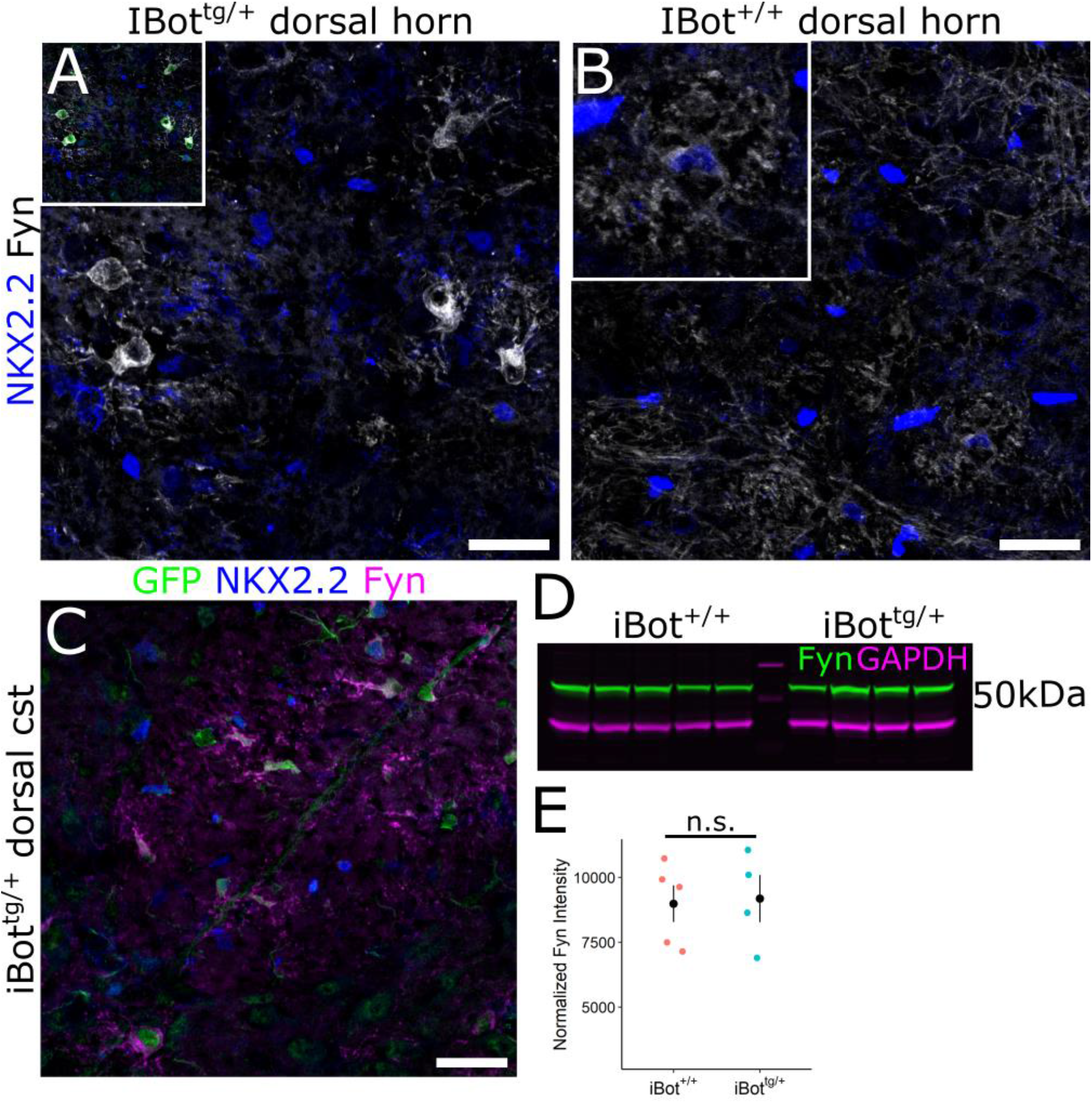
Fyn expression in iBot^tg/+^ spinal cords is concentrated in NKX2.2-BoNT/B-expressing OLs. **(A)** tg/+Immunolabeling for Fyn and NKX2.2 in iBot dorsal horn, inset shows overlay with GFP. **(B)** Immunolabeling for +/+ Fyn and NKX2.2 iBot dorsal horn, inset shows enlarged view of control premyelinating OL. **(C)** GFP, NKX2.2, tg/+ and Fyn labeling in iBot dorsal corticospinal tract. **(D)** Western blot for total Fyn (green) and GAPDH +/+ tg/+ (magenta) from P14-15 iBot and iBot spinal cord lysates. **(E)** Quantification of total Fyn signal normalized to +/+ 3 tg/+ 3 GAPDH from iBot (8.99×10 ± 705.49 a.u.) and iBot (9.18×10 ± 907.86 a.u.) spinal cord lysates (Welch two +/+ tg/+ sample t-test, t(6.06) = −0.168, p = 0.872, n = 5 iBot and 4 iBot mice). Scale bar = 25 μm **(A,B,C)**.

To determine whether the overall level of Fyn expression was enhanced in the spinal cords of iBot^tg/+^ mice, we performed Western blotting for Fyn using spinal cord lysates from iBot^+/+^ and iBot^tg/+^ mice (Figure 9D). Normalization to GAPDH indicated that, while not enhanced, overall levels of Fyn in the iBot^tg/+^ mice were very similar to controls (Figure 9E), despite the significant reduction in overall OL density. This can likely be explained, at least in part, by the increased prevalence of iBot^tg/+^ OLs in the premyelinating stage, when *Fyn* expression normally peaks. However, qualitatively, the level of Fyn staining associated with the iBot^tg/+^ OLs often appeared higher even than that of NKX2.2+ premyelinating OLs seen in control mice (Figure 9B, inset), suggesting that an overaccumulation of Fyn in the iBot^tg/+^ OLs may also contribute to the overall levels of Fyn detected in the iBot^tg/+^ spinal cord lysates.

## Discussion

Robust and timely production of new, myelinating OLs is essential for successful CNS myelination in development as well as efficient myelin remodeling in adulthood. Here we have shown that impairment of a subset of SNARE dependent processes, through toxin-mediated cleavage of VAMP2/3, resulted in a markedly diminished capacity to produce myelinating OLs in the developing mouse spinal cord, resulting in hypomyelination and significant motor impairment. Our data indicate that VAMP2/3 become critical to the development of OL lineage cells once they have exited cell cycle and begun to differentiate. Interestingly, the production of OPCs, as well as their capacity to self-renew and initiate differentiation, appeared largely unaffected by loss of intact VAMP2/3. During differentiation, however, OLs lacking intact VAMP2/3 appeared to stall in the premyelinating stage, ultimately failing to reach maturity. In addition to OL lineage cells, NG2-cre also drives expression of BoNT/B-LC in vascular pericytes. Labeling for Laminin+ endothelial cells in the iBot^tg/+^ mice, however, did not reveal any gross abnormalities in spinal cord vasculature that might otherwise account for the impaired maturation of OL lineage cells.

BoNT/B specifically cleaves VAMP1, 2, and 3, but not VAMP4, 5, 7, 8, or other SNARE proteins (Yamamoto et al., 2012). Since OL lineage cells primarily express VAMP2-4 and VAMP7 (Feldmann et al., 2009; Zhang et al., 2014), the observed effects are likely to have been mediated by cleavage of VAMP2/VAMP3. Although Cre is activated in vascular mural cells in addition to OL lineage cells in NG2-cre mice (Zhu et al., 2008), it is unlikely that the phenotype is due to off-target effects of by VAMP cleavage in non-OL lineage cells, as similar cell-autonomous myelination defects occur in other mouse lines that target BoNT/B-LC to OL lineage cells, such as iBot;Pdgfra-creER and iBot;Cnp-cre (Pan et al., BioRxiv; Lam et al., personal communication). In addition, a mouse protein BLAST search for the amino acid sequence flanking the BoNT-B cleavage site [GASQFE] (Schiavo et al., 2000; Binz et al., 2010) returned VAMP1, 2, and 3 as the only proteins exhibiting a complete match, further corroborating the interpretation that the effects observed in the iBot^tg/+^ mice are likely to have been caused by cleavage of VAMP2/3 in OL lineage cells.

It is interesting that we did not observe any compensatory response among OPCs to the deficiency in OL number or myelination. For comparison, jimpy and shiverer mice, two mutant mouse lines with a hypomyelination phenotype, both show enhanced proliferation and density of OPCs as a result of insufficient myelination (Roach et al., 1983; Nave et al., 1987; Wu et al., 2000; Bu et al., 2004). One possibility is that loss of VAMP2/3 impairs the ability of OPCs to respond to abnormal environmental conditions such as hypomyelination. Alternatively, the prevalence of premyelinating OLs, despite not exceeding the total number of OLs in control animals, could discourage the response of OPCs to hypomyelination through factors specifically expressed in the premyelinating stage. While these possibilities are speculative, they may be relevant in the context of demyelinating disease, where identification of factors that could desensitize OPCs to their environment may provide new targets for therapeutic strategies aimed at promoting remyelination.

Our data indicate that loss of VAMP2/3 function in the OL lineage primarily impacts cells in the early OL stage of development and suggest that VAMP2/3 dependent processes play a major role in managing this crucial period of OL differentiation. EdU pulse chase labeling of proliferative cells in developing iBot^tg/+^ and control mice indicated that the rate of new OL production was not significantly affected by loss of VAMP2/3, suggesting that VAMP2/3 function is not critical to the development of OL lineage cells prior to the onset of differentiation. This was further supported by the observation that the density of NKX2.2^high^ actively differentiating OLs in the dorsal column was similar between iBot^tg/+^ mice and controls. The reduced proportion of OLs labeled by the mature OL markers GSTπ and ASPA in the iBot^tg/+^ mice suggested that their diminished capacity to produce OLs may have been due to a defect in maturation of differentiating OLs. Immunolabeling for NKX2.2 in the dorsal column further indicated that the reduced NKX2.2+ OL density was specific to a lack of fully mature OLs, which re-express NKX2.2 at low levels. Together, this suggests that VAMP2/3 serves an essential function late in the OL differentiation program, possibly by mediating the secretion and/or trafficking of a hypothetical molecule X (see Figure 10) that is required for OLs to become fully mature. Mice in which expression of BoNT/B is regulated by the Cnp promoter are defective in elaborating myelin membranes (Lam, Zuchero, et al., personal communication). In addition, the increased prevalence of iBot-expressing OLs with premyelinating morphology indicated that their progression through this stage may be extended, or that they may linger in the final stages of maturation, prior to being lost.

**Figure 10:**
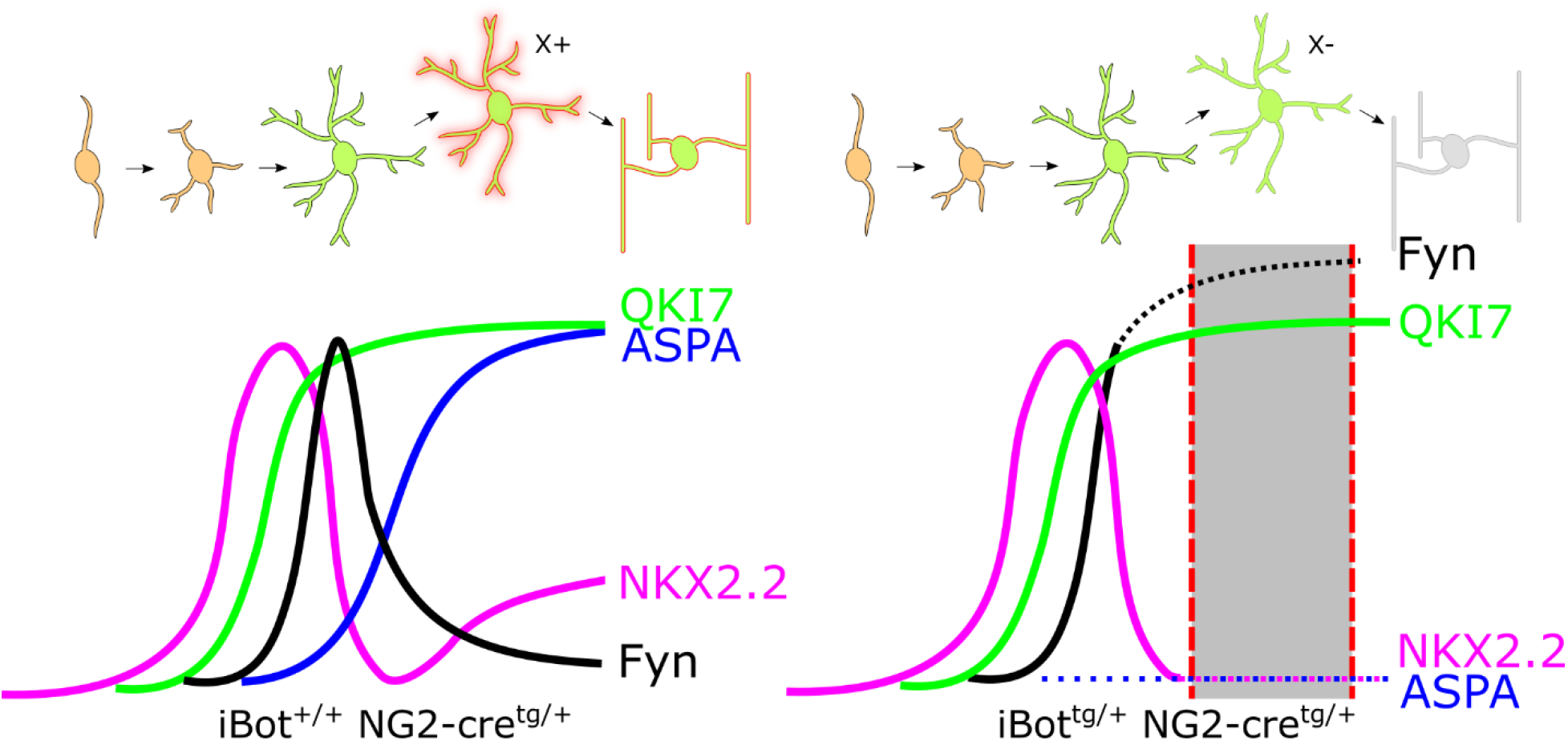
Proposed model of OL lineage development and gene expression timing in iBot^+/+^ and iBot^tg/+^ spinal cords. Dotted lines indicate abnormal expression timing observed in the iBot^tg/+^ mice relative to control, while the shaded region denotes the approximate stage in which iBot^tg/+^ OLs are lost. Impaired VAMP2/3 dependent secretion and/or trafficking of a hypothetical molecule X is indicated at the transition between premyelinating and myelinating stages of development.

Given that the rate of new OL production did not appear affected in the iBot^tg/+^ mice, the reduced overall density of QKI7+ OLs and the strong reduction in the density of mature NKX2.2^low^ re-expressing OLs implies that VAMP2/3-defecient OLs ultimately undergo cell death. Staining for cCasp3 indicated that, while not severe, the iBot^tg/+^ mice do exhibit increased cell death in the dorsal column, largely occurring within OL lineage cells labeled by Olig2. One possibility is that delayed development, or a prolonged premyelinating stage, may trigger apoptosis driven by an intrinsic cell death timer. It has been shown that the premyelinating stage of OL development is under strict temporal control (Czopka et al., 2013; Hill et al., 2014) and that a significant portion of newly generated OLs ultimately undergo apoptosis (Barres et al., 1992; Trapp et al., 1997; Hughes et al., 2018). Therefore, it is possible that a prolonged early OL stage in VAMP2/3-deficient OLs could trigger apoptosis by impairing timely axon-OL contact and initiation of myelination.

While our results suggest many of the iBot^tg/+^ OLs seem to be stalled in the premyelinating stage, some did appear to display morphology consistent with actively myelinating cells. This may indicate that VAMP2/3 function becomes critical just as premyelinating OLs begin to contact axons and initiate myelination. As a result, iBot^tg/+^ OLs observed with a more mature morphology may have been still in the initial stages of myelination, prior to undergoing cell death. Alternatively, there may be heterogeneity among OLs in their requirement for VAMP2/3 to complete differentiation successfully. This heterogeneity could either be innate and cell-specific or determined by the microenvironment around a given OL, including the type and number of axons contacted or available for contact. It is also possible, however, that those iBot^tg/+^ OLs which achieve a more mature developmental stage are derived from escaped OPCs expressing EGFP but not BoNT/B-LC or cells where the dosage of BoNT/B-LC produced is insufficient to fully prevent their development.

Our data suggest that a VAMP2/3 dependent mechanism regulates the successful transition from the premyelinating to myelinating stage of OL development. It is possible that this mechanism involves the SNARE mediated trafficking and/or secretion of some molecule X that is required for differentiating OLs to progress beyond the premyelinating stage, and that functional loss of this molecule X in the iBot^tg/+^ OLs is what leads to their stalled differentiation (Figure 10). Another indication that the iBot^tg/+^ OLs undergo an extended premyelinating stage is their apparent over-expression of Fyn, which may accumulate due to their stalling in a normally high Fyn-expressing stage. Our immunofluorescent localization of Fyn in iBot^tg/+^ premyelinating OLs showed that Fyn appeared to be highly expressed and present throughout both the soma and distal processes. This indicated that, in addition to being expressed and exhibiting markers of activation (SFK pY416+), Fyn was also being successfully trafficked out to sites of potential axon-OL contact, suggesting that loss of VAMP2/3 does not directly impair these processes.

Western blotting for Fyn indicated that the overall level of Fyn expressed in iBot^tg/+^ spinal cords was similar to controls, despite having a significantly reduced OL density. As *Fyn* expression typically peaks in the premyelinating stage, the increased prevalence of premyelinating OLs in iBot^tg/+^ mice would be expected to offset the reduction in total Fyn levels to a degree. However, immunofluorescent labeling for Fyn further suggested that the iBot^tg/+^ OLs may exhibit even higher levels of Fyn than would be normally expected from peak *Fyn* expression during the premyelinating stage. In addition, we observed that the iBot^tg/+^ OLs displaying the strongest Fyn immunoreactivity were largely NKX2.2-, indicating they had completed the initial downregulation of NKX2.2 that occurs in the final stages of maturation. This provided further support for our interpretation that the increased level of Fyn seen in the iBot^tg/+^ OLs may have resulted, at least in part, from a prolonged accumulation of Fyn due to their stalled progression in the premyelinating stage, as summarized in Figure 10. Additionally, negative feedback on Fyn expression that might normally be exerted by mature OLs would be greatly reduced in the iBot^tg/+^ mice, which may further drive over-expression of Fyn. Alternatively, it is also possible that prolonged Fyn signaling itself could prevent differentiating OLs from reaching full maturity due to a failure to downregulate Fyn via a VAMP2/3 dependent mechanism. While it is unclear based on our data if VAMP2/3 is required for the downregulation of *Fyn* following differentiation, it is possible that the mechanism responsible is impaired by loss of VAMP2/3, ultimately leading to sustained Fyn expression and undermining OL lineage cell development.

Overall, our results show that VAMP2/3 play an essential role in regulating spinal cord OL differentiation during development. Differentiating OLs lacking functional VAMP2/3 appeared to undergo an extended, and ultimately unsuccessful, premyelinating stage, exhibiting expression profiles that indicated they were neither actively differentiating nor fully mature. Conversely, OPCs appeared relatively insensitive to loss of VAMP2/3 and maintained their capacity for self-renewal as well as their ability to initiate differentiation. Taken together, our data suggest that a VAMP2/3 dependent mechanism active late in the differentiation process, during the transient downregulation of NKX2.2, is required for successful maturation of differentiating OLs to occur. While the precise nature of this mechanism is unclear, it may involve the trafficking, secretion, or membrane presentation of key axon-OL signaling factors via a VAMP2/3 specific pathway.

## Acknowledgements

This work was supported by National Institutes of Health grants R01NS116182 and R01NS073425 and National Multiple Sclerosis Society grants PP-1809-32554 and RG-1612-26501 to Akiko Nishiyama. The Leica SP8 confocal was purchased using funds from NIH Instrumentation Grant S10 OD016435 to Akiko Nishiyama. We thank Youfen Sun for maintaining transgenic mouse lines, Dr. Chris O’Connell for assistance with confocal microscopy, as well as Jessica Ulbrich and Owen Greene for assistance with immunofluorescent labeling.

---

**Video 1:** Example video of a representative iBot^tg/+^ mouse at P13 demonstrating severely impaired coordination and motor function, followed by a control P13 mouse for comparison starting at 27 seconds.

